# Pilot identification of the Live-1/Prox-1 expressing lymphatic vessels and lymphatic elements in the unaffected and affected human brain

**DOI:** 10.1101/2021.09.05.458990

**Authors:** O. Semyachkina-Glushkovskaya, I. Fedosov, N. Navolokin, A. Shirokov, G. Maslyakova, A. Bucharskaya, I. Blokhina, A. Terskov, A. Khorovodov, D. Postnov, J. Kurths

## Abstract

We report here a pilot identification of the presence of the lumenized Lyve-1/Prox-1-expressing vessels with distinct walls composed of a single endothelial layer in the unaffected brain and with intraventricular hemorrhages. These lymphatic vessels (LVs) have valves and an undulating shape in the distal region that is the classical characteristic of lymphatic precollectors. Furthermore, we identified Lyve-1/Prox-1-expressing lymphatic elements in the enlarged perivascular spaces. These pioneering results might be a revolutionary step in our understanding of anatomy and physiology of the cerebral lymphatics and stimulate a reassessment of basic assumptions in the aetiology of brain diseases associated with lymphatic dysfunction. The discovery of the cerebral LVs is a crucial step in the development of breakthrough technologies of modulations of lymphatic removing of toxins and blood from the brain.

**One Sentence Summary:** Lyve1/Prox1-expressing lymphatic vessels and elements in the unaffected and affected human brain.

---

The classical characteristics of the central nervous system (CNS) is a lack of the lymphatic vessels (LVs). However, it has become more and more evident that the CNS is not a passive “immune-privileged” organ, but it is rather a compartment with highly active immune responses (1-3). Although the brain lacks “classical” lymphatics, tertiary lymphoid organs (TLOs) have been observed in the CNS of patients with various neurological diseases (2,4,5). The TLOs are lymph-node like structures which develop at the sides of chronic inflammation and play an important role in the interaction between the inflamed brain vasculature and the lymphatic proteins (2). The presence of LVs is one of the characteristics feature of TLOs in the peripheral tissues (4,5). However, it is still not clear whether LVs are present in TLOs of the inflamed brain. In a recent immunohistochemical study, lymphatic elements (LEs) have been explored in ten postmortem human brains (with and without neurological pathology) (6). This elegant study has identified LEs expressing proteins, such as the lymphatic vessel endothelial hyaluronan receptor 1 (Lyve-1) and podoplanin but lymphatic lumenized vessel-like structures were not found.

The LVs have been discovered in the dura mater of humans, rodents and zebrafish (3,7-11). However, the dura mater is not a component of the CNS, but covers, along with the other meningeal layers, both the brain and the spinal cord. For the first time, the presence of lymphocyte-containing channels in the CNS was described by Prineas in 1979 (12). He examined five human brains with various neurological diseases and observed thin-walled channels, which were indistinguishable from lymphatic capillaries in the peripheral tissues. However, Prineas identified the lymphatic channels using electron microscopy without the specific markers of lymphatic endothelium cells (LEC). This makes these findings debatable and the crucial question of whether LVs exist in the human brain remains open.

To address this problem, we examine here the affected brains of patients with primary and secondary intraventricular hemorrhages (IVH, n=34, Table S1) and in the unaffected brain (the control group, n=8, Table S2) of patients with a congestive heart failure and the development of pulmonary edema using immunostaining with two classical markers of LEC, such as the surface marker Lyve-1 and the nuclear transcription factor, such as the prospero homeobox protein 1 (Prox-1). We discuss in the Supplementary the benefits of using of these two LEC markers but also significant limitations of other LEC markers for the study of human brain, including particularities of human brain tissues, biopsy sampling and the rules of certification of brain death. We were focused on the study of the cerebral cortex, which is covered by three layers of the meninges with a network of conventional LVs (3,7-11) (Figure 1a). For the sake of consistency and reproducibility, in this pilot step we analyzed two regions of the human brain in the postcentral gyrus of the parietal lobe and in the inferior occipital gyrus. There is strong evidence that the meningeal lymphatic vessels (MLVs) play an important role in drainage and clearing of the brain tissues (3,13,14). If so, it is logical to assume the presence of LVs in the cerebral cortex that can explain how fluids drain from the brain tissues to MLVs. However, LVs have not yet been discovered directly in the human and animal brain parenchyma. Therefore, here we aimed to study, whether LVs and LEs are present in the mature human brain.

**Fig. 1.**
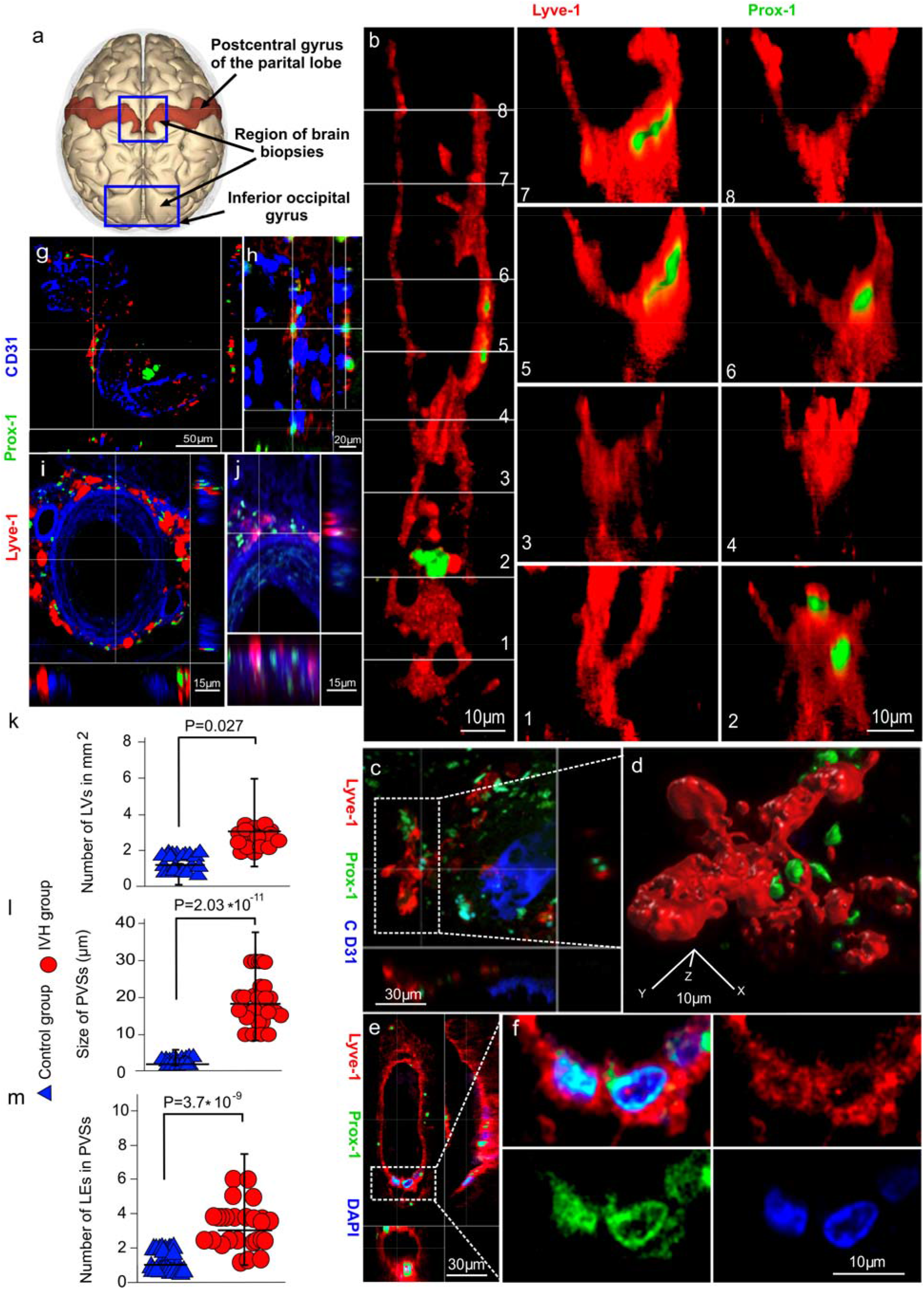
The pilot identification of LVs and LEs in the human brain. **a**, Schematic illustration of region of brain biopsies; **b**, Representative images of the lumenized Lyve-1/Prox-1-expressing vessel with distinct walls composed of a single endothelial layer in coronal projection: (1 and 2) – the undulating shape in distal region of LVs, (3 and 4) – valve in LV, (5-8) – the walls of LV and colocolization of Lyve1 with Prox1; **c** and **d**, 3D reconstruction of a region of interest demonstrating the initial lymphatics with the expression of Lyve-1 and Prox-1; **e**, Lyve-1/Prox-1 expressing LVs in axial projection; **f**, Colocolization of Prox-1 with DAPI and Lyve-1 in the lumenized Lyve-1/Prox-1-expressing vessel, **g-j**, Representative images of LEs expressing Lyve-1 and Prox-1 around CD31 vessels in sagittal and axial projections; **k**, Quantifications of the number of LVs in mm^2^ of the brain tissue; **l** and **m**, quantifications of size of PVSs and the number of LEs in PVSs; **k-m**, each dot represents a single value combined in one group, n=3 for the control group and n=5 for the IVH group, Me Q [25-75], two-tailed Mann–Whitney U test.

Figures 1 b and e demonstrate the lumenized Lyve-1/Prox-1-expressing vessels and clearly show distinct walls composed of a single endothelial layer. We found valves and a classical undulating shape in the distal region of Lyve-1/Prox-1 vessels (Fig. 1b(_1-4_), Videos 1-3). The LVs were found only in the inferior occipital gyrus. The colocalization with two LEC markers clearly revealed that an expression of the main LEC transcription factor, Prox-1, was indeed delectable with the Lyve-1 protein and with DAPI (Fig. 1b(_2,5,6,7_),d and f, Videos 4 and 5). The videos 1,3,4 show the Lyve-1/Prox-1 vessels in 3D projections. Furthermore, we identified Lyve-1/Prox-1-expressing LEs in 3 of 8 brains in the control group, in 8 of 34 brains of patients with IVH (Fig. 1 g-j, Tables S1 and S2). Among the cases with obtained LEs, LVs were found in 3 unaffected brain and in 5 of 8 brains with IVH. The number of LVs was higher in the IVH group vs. the control group (Fig. 1 k). The diameter of LVs was not evaluated due to the limited number of observed LVs in the tested brain.

The immunohistochemical (IHC) assay of brain slices stained for CD31 and Lyve1 revealed that the Lyve1+structures were identified along CD31+ capillaries, arterioles and venules with an enlarged perivascular space (PVSs) (Fig. S1, Table S3). The size of PVSs and the average the number of Lyve1+ structures in PVSs were much higher in the IVH group compared with the control group (Figure 1 l and m, Table S3). There were no differences between both two groups in the diameter of LEs (Table S3). The LEs were found in both the postcentral gyrus of the parietal lobe and in the inferior occipital gyrus.

The LVs can be distinguished from blood vessels due to the presence of Lyve-1 (15). However, the Lyve1-positive macrophages are present in the lymphatic and non-lymphatic regions in the meninges of rats (16). We therefore examined the brain slices immunostained for CD68 (macrophages), CD31 (the blood endothelial cell marker) and Lyve-1 (LEC marker) to rule out the possibility of artifacts caused by Lyve-1. Our confocal and IHC analysis revealed that the CD68+ cells are not part of Lyve-1+ structures, but they are expressed close to each other along CD31+ vessels (Fig S2, S3, and Video 6). We also found the presence of CD68 cells directly in the lumen of Lyve-1-structures (Fig. S2b).

Collectively, our pilot results clearly demonstrate the presence of lumenized Lyve-1/Prox-1-expressing vessels with distinct walls composed of a single endothelial layer. These LVs have valves and an undulating shape in the distal region that is a classical characteristic of a lymphatic precollectors (Table S4). Furthermore, we identified Lyve-1/Prox-1-expressing LEs in the healthy human brain and with IVH.

The presence of LVs in the human brain is a missing crucial link in the concept of the cerebral lymphatic system, which warrants a substantial revision of our knowledge about the role of the cerebral LVs in the brain pathology. The lymphatic pathway of removing of cells and macromolecules from the brain has been shown in many studies (3,8,13,14,17) and the cerebral LVs can be an initial pathway of clearance of toxins and metabolites from the brain that conceptually improves the glymphatic theory (18,19). The cerebral LVs can be a key player in the communications between the subarachnoid space (SAS) and the brain parenchyma creating the drainage tunnel for the interstitial fluid outflow into SAS that is described in our mathematical model of brain lymphatic drainage (Fig. S5, S6, Video S7).

To conclude, further studies of anatomy and functions of cerebral LVs, including traffic of immune cells, will be a revolutionary step in a reassessment of basic assumptions in neuroimmunology and the aetiology of brain diseases associated with a lymphatic system dysfunction. The discovery of the cerebral LVs is a crucial step in the development of breakthrough technologies of therapeutic modulation of lymphatic removing of blood and toxins from the brain that we revealed in our study on mice with a model of intraventricular hemorrhages (20).

## Acknowledgements

We thank Saranceva Elena and Elmira Kaybeleva for the graph design and preparation of video S7.

## Author contributions

Conceptualization: O.S-G.

Methodology: I.F., N.N., A.Sh., A.B., G.M.

Investigation: O S.G., I.F., N.N., A Sh., I.B., A.T., A.K.

Visualization: A.Sh. and I.F.

Project administration: O.S.-G. and J.K.

Supervision: O S-G. and J.K.

Writing – original draft: O S-G.

Writing – review & editing: J.K. and O.S-G.

## Data and materials availability

All data are available in the manuscript or the Supplementary.

