## Supplemental material for "Pilot identification of the Live-1/Prox-1 expressing lymphatic vessels and lymphatic elements in the unaffected and affected human brain"

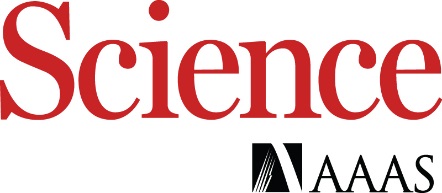

Supplementary Materials for

Pilot identification of the Live-1/Prox-1 expressing lymphatic vessels and lymphatic elements in the unaffected and affected human brain

O. Semyachkina-Glushkovskaya^1,2*,^ I. Fedosov^2^, N. Navolokin^2,3^, A. Shirokov^2,4^, G. Maslayakova^2,3^, A. Bucharskaya^2,3^, I. Blokhina^2^, A. Terskov^2^, A. Khorovodov^2^, D. Postnov^2^, J. Kurths^1,2,5^.

**This PDF file includes:**

Materials and Methods

Supplementary Text

Figs. S1 to S6

Tables S1 to S4

References (25-29)

Captions for Movies S1 to S7

**Other Supplementary Materials for this manuscript include the following:**

**Movies S1 to S7**

Movie S1.

Representative image of 3D reconstruction of the lumenized Lyve-1/Prox-1-expressing vessel with distinct walls composed of a single endothelial layer in coronal, sagittal and axial projections. Lyve-1 (red) and Prox-1 (green).

Movie S2.

Animated sequence of cross-sections of undulating shape in the distal region of Lyve-1/Prox-1-expressing vessel. Lyve-1 (red) and Prox-1 (green).

Movie S3.

3D representative image of undulating shape of the Lyve-1/Prox-1-expressing vessel. Lyve-1 (red) and Prox-1 (green).

Movie S4.

Representative image of 3D reconstruction of the lumenized Lyve1/Prox1-expressing vessel and colocolization of Lyve-1 with Prox-1 and Prox-1 with DAPI in coronal, sagittal and axial projections. Lyve-1 (red), Prox-1 (green), DAPI (blue).

Movie S5.

Colocolization of Prox-1 with DAPI in coronal, sagittal and axial projections. Prox-1 (green), DAPI (blue).

Movie S6.

Representative image of 3D reconstruction of Lyve-1-expressing lymphatic elements around CD31 blood vessel and CD68-expressing cells. Lyve-1 (red), CD68 (green), CD31 (blue).

Movie S7.

The schematic representation of LVs in the human brain and a model of brain lymphatic drainage.

**Data S1 to S4**

**Table S1**. The group of patients, who died from the cerebral hemorrhage.

**Table S2**. The control group of patients, who died from the congestive heart failure with the development of pulmonary edema.

**Table S3**. The quantitative analysis of PVS, CD31, CD68 and Lyve-1 positive structures.

**Table S4**. The general characteristics of the initial lymphatics and precollectors.

Materials and Methods

**Human samples**. Autopsy specimens of human brains and meninges were obtained from the Department of Pathological Anatomy at the Saratov State Medical University (average age 65). All brain samples were obtained within the first 2-3 hours after death, immediately fixed and stored in a 10% formalin solution. The instigations were performed on the following groups: 1) the control group included the unaffected brain obtained from patients died from the congestive heart failure and the development of pulmonary edema (n=8); 2) patients died after primary and secondary intraventricular hemorrhages (IVH), n=34 (Tables S1 and S2).

**Immunohistochemistry of human brains**

For identification of the lymphatic vessels (LVs) and the lymphatic elements (LEs), we used two classical markers of the lymphatic endothelium, such as the lymphatic vessel endothelial hyaluronan receptor 1 (Lyve-1, the surface marker) and the prospero homeobox 1 protein (Prox-1, the nuclear marker). Notice that, there are no ideal markers of the lymphatic endothelial cells (LEC) in the human brain. Furthermore, the human brain samples cannot be taken immediately as in animals due to the rules of the certification of brain death that requires minimum 2-3 hours for confirmation of brain death (1). However, studies in both humans and experimental animals have shown that oxygen stores, global electrical activity, glucose, ATP stores and expression of many proteins in the brain are lost within minutes of interrupted blood flow (2,3). Additionally, the marker of LEC, such as chemokine (C-C motif) ligand 21 (CCL-21) is not expressed in the human brain (4). The other marker of LEC, vascular endothelial growth factor receptor 3 is widely distributed in the human brain tissues and cannot be specific marker of LEC (5-9). Mezey et al. demonstrated expression of podoplanin, which is the integral membrane protein of LEC, only in the human meninges and identified LEs in the parietal and frontal cortex of human brain using one LEC marker, such as Lyve-1 (5). Blood vascular endothelial cells express a pan endothelial cell marker (CD31) but not Prox-1 (10). However, LVs express Prox-1 and can also express CD31 (10). Thus, the general opinion of study of LVs together with CD31 cannot be ideal approach. The LVs can be distinguished from blood vessels due to the presence of Lyve1 (11). However, Lyve-1+ cells are also observed in intra-embryonic arterial and venous endothelium (6). The development of lymphatic structures starts with the up-regulation of the lymphatic marker Prox1 in a subset of venous endothelial cells (12,13). Thus, Lyve-1 and Prox-1 looks as more promising and reliable markers of LVs but only for the mature brain. Therefore, we identified LVs in the mature brains using Lyve-1 and Prox-1 with cell surface and nuclear localization, respectively. For the sake of consistency and reproducibility, in this pilot step we analyzed two regions of the human brain in the postcentral gyrus of the parietal lobe and in the inferior occipital gyrus.

For confocal visualization of LVs and LEs in the human brains, the protocol for the immunohistochemical (IHC) analysis was used with two markers of the lymphatic endothelial cells, such as Lyve-1 (ab219556; Abcam, Biomedical Campus Cambridge, Cambridge, UK, 1:500) and Prox-1 (NBP1-30045; Novus Biologicals, Littleton, USA, 1:500), for the marker of blood endothelium CD31 (ab187377; Abcam, Biomedical Campus Cambridge, Cambridge, UK, 1:500) and for marker of macrophages CD68 (ab 201973, Abcam, Biomedical Campus Cambridge, Cambridge, UK, 1:500), DAPI for stain dsDNA (ab228549; Abcam, Biomedical Campus Cambridge, Cambridge, UK).

For the ICH analysis, brain tissues were collected and free-floating sections were prepared. Pieces of the brain, measuring 2x2 сm, were fixed for 48 hours in a 4% saline solution-buffered formalin, then sections of the brain with a thickness of 40-50 microns were cut on a vibrotome (Leica Microsystems GmbH, Germany). Brain sections were processed according to the standard IHC protocol with the corresponding primary and secondary antibodies. Confocal microscopy of human brain sections was performed using a Leica SP8 confocal laser scanning microscope (Leica Microsystems GmbH, Germany). The nonspecific activity was blocked by 2-hour incubation at room temperature with 10% BSA in a solution of 0.2% Triton X-100 in PBS. Solubilization of cell membranes was carried out during 1-hour incubation at room temperature in a solution of 1% Triton X-100 in PBS. Incubation with primary antibodies in a 1:500 dilution was performed overnight at 4 ° C: with mouse antibodies to Prox1 (1:500; NBP1-30045; Novus Biologicals, Littleton, USA); rabbit antibodies to Lyve-1 (1:500; ab219556; Abcam, Biomedical Campus Cambridge, Cambridge, UK); mouse antibodies to CD31 (1:500; ab187377; Abcam, Biomedical Campus Cambridge, Cambridge, UK) and mouse antibodies to CD68 (1:500; ab201973; Abcam, Biomedical Campus Cambridge, Cambridge, UK), DAPI (ab228549; Abcam, Biomedical Campus Cambridge, Cambridge, UK). At all stages, the samples were washed 3-4 times with 5-minute incubation in a washing solution. Afterward, the corresponding secondary antibodies were applied (goat anti-mouse IgG (H+L) Alexa Four 488; goat anti-mouse IgG (H+L) Alexa Four 555 and goat anti - rabbit IgG (H+L) Alexa Four 647; Invitrogen, Molecular Samples, Eugene, Oregon, USA). At the final stage, the sections were transferred to the glass and 15 µl of mounting liquid (50% glycerin in PBS with DAPI at a concentration of 2 µg/ml) was applied to the section, then the preparation was covered with a cover glass and confocal microscopy was performed.

IHC examination was performed on paraffin sections, using a standard double-immunohistostaining protocol with monoclonal antibodies to Lyve1 (ab219556, Abcam, Biomedical Campus Cambridge, Cambridge, Abcam, UK, 1:500), CD-31 (ab187377, Abcam, Biomedical Campus Cambridge, Cambridge, Abcam, UK, 1:500), CD-68 (ab201973, Abcam, Biomedical Campus Cambridge, Cambridge, Abcam, UK, 1:500).

Initially, endogenous peroxidase was blocked by adding 0.3% hydrogen peroxide to the sections for 10 min followed by washing the sections in Phosphate-Buffered Saline (PBS). The antigen retrieval was conducted using a microwave oven in a Trilogy Buffer, nonspecific background staining was blocked in PBS containing 0.5% BSA and 0.5% casein for 10 min, after which the sections were washed in PBS for 5 min. To distinguish blood vessels from LVs, we used a double-staining protocol. The sections were incubated with first antibody - diluted mouse anti-CD31 antibody (ab187377, Abcam, UK, 1:500) for 1 h at room temperature, afterward, the sections were washed in PBS. The immunohistochemical reaction was visualized with HiDef. Detection HPR Polymer System (Cell Marque, USA) with DAB chromogen visualization (brown color). In the second step, we used rabbit anti-Lyve-1 antibody (ab219556, UK, 1:500 ) with AEC chromogene visualization (red color). Then the sections counterstained with hematoxylin for 1 min, washed in water, and finally embedded into aqueus mounting medium.

To distinguish macrophages from LVs, we used a similar double-staining protocol. In the first step, we used diluted rabbit anti-CD68 antibody (ab201973, Abcam, UK, 1:500) with DAB chromogen visualization (brown color). In the second step, we used anti-Lyve-1 antibody with AEC chromogene visualization (red color).

The morphological studies were performed in at least 10 fields of view of each brain sample. The morphometric measurements of the size of blood vessel, the size of PVS, the number and diameter of Lyve-1 positive LVs, and the number of macrophages were performed using a µVizo-103 Medical Microvizor (LOMO, Russia) and magnification 246.4, 774.0. Morphometric measurements were performed only in brain samples with positive expression of Lyve-1. The diameter of Lyve-1-positive LVs was measured along CD31-positive microvessels (up to 100 µm).

**Image analysis.**

Multichannel image stacks were acquired with a Leica TCS SP8 confocal system (Leica Microsystems GmbH, Germany) with 20×0.7 dry objective lens. Images post processing was performed with FIJI software (NIH). It included median filtration; brightness and contrast correction; and crop of original stacks. 3D visualizations of image stacks were performed with Vaa3D software (25).

**Statistical analysis**

All statistical analysis was performed using Microsoft Office Excel and SPSS 17.0 for Windows software. Normality of group distribution was checked using the Shapiro-Wilk criterion. For nonparametric data distribution, significance of differences between specific groups was determined using Mann-Whitney criterion (U/Z test) with calculation of median (Me), 25th and 75th percentiles Q (25-75), maximum and minimum. In this method, median differences were determined at Z≥1.96 at a significance level of p<0.05 (with more than 95% probability). No statistical methods were used to predetermine sample size.

Table S1. The group of patients, who died from the cerebral hemorrhage

| **N** | **HSB Sample no** | **Age** | **Sex** | **Structure** | **Neuropathology Macroscopic description** | **Neuropathology Microscopic description** | **Clinical diagnosis** | **Cause of death** | **Postmortem time, h** |
| --- | --- | --- | --- | --- | --- | --- | --- | --- | --- |
| 1 | 446/19 | 52 | F | Cerebral cortex (Frontal lobe). | In the area of the temporoparietal lobe of the right hemisphere, a cavity measuring 7x5x4.5 cm filled with a blood clot, with a defect in the wall, through which blood flows into the lumen of the ventricles. | Perivascular and pericellular edema, fresh hemorrhage in the brain matter | Intracerebral hemorrhage with formation of intracerebral hematoma in the right hemisphere and rupture into the ventricular system of the brain. Left-sided hemiplegia. | The cerebral edema with occlusion of the cerebellar tonsils into the greater occipital foramen. | 3.0 |
| 2 | 465/19 | 72 | M | Cerebral cortex (Temporal lobe) | In the basal nuclei of the right hemisphere there is a cavity of 5x4x3 cm filled with blood clot, there is a defect in the wall, through which blood enters the ventricular lumen. | Perivascular and pericellular edema, fresh hemorrhage in the brain matter.  **Immunohistochemical staining of cerebral cortex tissues revealed Lyve1/Prox-1 positive elements.** | Intracerebral hemorrhage in the right hemisphere with formation of intracerebral hematoma. Dysarthria, left-sided hemiparesis up to hand plegia. | The cerebral edema with occlusion of the cerebellar tonsils into the greater occipital foramen. | 2.0 |
| 3 | 498/19 | 72 | F | Cerebral cortex (Frontal lobe). | In the area of the temporoparietal lobe of the left hemisphere, a cavity measuring 7x5x4.5 cm filled with a blood clot, with a defect in the wall, through which blood flows into the lumen of the ventricles. | Perivascular and pericellular edema, fresh hemorrhage in the brain matter. | Parenchymatous ventricular hemorrhage with formation of intracerebral hematoma in the left cerebral hemisphere with blood rupture into the brain ventricles, motor-sensory aphasia, right-sided hemiplegia. | The cerebral edema with occlusion of the cerebellar tonsils into the greater occipital foramen. | 2.1 |
| 4 | 11/20 | 79 | F | Cerebral cortex (Parietal lobe). | There is blood in the lumen of the fourth and lateral ventricles. | Perivascular and pericellular edema, fresh hemorrhage in the brain matter.  **Immunohistochemical staining of cerebral cortex tissues revealed Lyve1/Prox-1 positive elements.** | Spontaneous recurrent intracerebral hemorrhage of multiple localization. | The cerebral edema with occlusion of the cerebellar tonsils into the greater occipital foramen. | 2.0 |
| 5 | 10/20 | 65 | F | Cerebral cortex (Parietal lobe). | In the right occipital lobe, a softening foci of smeared consistency with indistinguishable structure, grayish-red in color and 3x4x4 in size, blood in the lumen of the left lateral ventricle. | Perivascular and pericellular edema, fresh hemorrhage in the brain matter  **Immunohistochemical staining of cerebral cortex tissues revealed Lyve1/Prox-1 positive elements.** | Parenchymatous ventricular hemorrhage with formation of intracerebral hematoma in the left cerebral hemisphere with blood rupture into the ventricles. | The cerebral edema with occlusion of the cerebellar tonsils into the greater occipital foramen. | 2.2 |
| 6 | 76/20 | 76 | F | Cerebral cortex (Occipital lobe). | In the left occipital lobe, a softening foci of smeared consistency with indistinguishable structure, grayish-red in color and 3x4x3 in size, blood in the lumen of the left lateral ventricle. | Perivascular and pericellular edema, fresh hemorrhage in the brain matter. | Parenchymatous-ventricular hemorrhage in the left cerebral hemisphere with formation of subdural and intracerebral hematomas with blood rupture into ventricles and subarachnoid space. | The cerebral edema with occlusion of the cerebellar tonsils into the greater occipital foramen. | 2.4 |
| 7 | 41/20 | 71 | M | Cerebral cortex (Parietal lobe). | In the area of the temporoparietal lobe of the left hemisphere there is a 7x5 cm cavity filled with blood clot, there is a defect in the wall, through which blood flows into the lumen of the ventricles. | Perivascular and pericellular edema, fresh hemorrhage in the brain matter.  **Immunohistochemical staining of cerebral cortex tissues revealed Lyve1/Prox-1 positive vessels and elements**. | Parenchymatous ventricular hemorrhage with formation of intracerebral hematoma in the left cerebral hemisphere with blood rupture into the brain ventricles, motor-sensory  aphasia, right hemiplegia. | The cerebral edema with occlusion of the cerebellar tonsils into the greater occipital foramen. | 2.0 |
| 8 | 39/20 | 67 | M | Cerebral cortex (Parietal lobe). | In the area of basal nuclei there was a softening foci of smeary consistency with indistinguishable structure, grayish-red in color and 3*4*3 in size, there was blood in the ventricular lumen. | Perivascular and pericellular edema, fresh hemorrhage in the brain matter  **Immunohistochemical staining of cerebral cortex tissues revealed Lyve1/Prox-1 positive vessels and elements.** | Consequences of intracerebral hemorrhage in the left middle cerebral artery basin with formation of intracerebral hematoma in the basal ganglia with rupture into the brain ventricles. | The cerebral edema with occlusion of the cerebellar tonsils into the greater occipital foramen. | 3.0 |
| 9 | 173/20 | 75 | F | Cerebral cortex (Parietal lobe). | Hemorrhage in the basal nuclei of the left hemisphere with blood rupture into the lateral ventricles of the brain. | Perivascular and pericellular edema, fresh hemorrhage in the brain matter.  **Immunohistochemical staining of cerebral cortex tissues revealed Lyve1/Prox-1 positive vessels and elements.** | Parenchymatous ventricular hemorrhage in the left cerebral hemisphere with formation of intracerebral hematoma with blood rupture into the ventricles | The cerebral edema with occlusion of the cerebellar tonsils into the greater occipital foramen | 2.0 |
| 10 | 201/20 | 38 | M | Cerebral cortex (Temporal lobe). | In the left posterior connective artery there is a 0.5 cm diameter sacciform aneurysm with rupture. The aneurysm wall with a laminated gray-cross-colored thrombus. There is blood in the lumen of the 4th ventricle and lateral ventricles. | Perivascular and pericellular edema, fresh hemorrhage in the brain matter. | Massive parenchymatous ventricular hemorrhage with formation of intracerebral hematoma in the left cerebral hemisphere with blood rupture into the brain ventricles. | The cerebral edema with occlusion of the cerebellar tonsils into the greater occipital foramen. | 2.3 |
| 11 | 208/20 | 46 | M | Cerebral cortex (Parietal lobe). | In the subdural space on the right, there is a small amount of blood soaking the cerebral cortex. | Perivascular and pericellular edema, fresh hemorrhage in the brain matter.  **Immunohistochemical staining of cerebral cortex tissues revealed Lyve1/Prox-1 positive elements.** | Spontaneous subarachnoid hemorrhage with the formation of a subdural hematoma in the right hemisphere of the brain. | The cerebral edema with occlusion of the cerebellar tonsils into the greater occipital foramen | 2.1 |
| 12 | 362/20 | 53 | F | Cerebral cortex. | Hemorrhage 6x3 cm in the parietal lobe of the right cerebral hemisphere | Perivascular and pericellular edema, fresh hemorrhage in the brain matter. | Parenchymatous hemorrhage with formation of intracerebral hematoma in the right cerebral hemisphere. | The cerebral edema with occlusion of the cerebellar tonsils into the greater occipital foramen. | 2.5 |
| 13 | 383/20 | 67 | F | Cerebral cortex. | There was hemorrhage in the left brainstem nuclei with rupture into the fourth and lateral ventricles. | Perivascular and pericellular edema, fresh hemorrhage in the brain matter. | Parenchymatous ventricular hemorrhage in the left cerebral hemisphere with formation of intracerebral hematoma and blood rupture into the ventricles. | The cerebral edema with occlusion of the cerebellar tonsils into the greater occipital foramen. | 2.0 |
| 14 | 557/20 | 43 | M | Cerebral cortex. | In the right cerebral hemisphere from the frontal to the parietal lobes there is a softening foci of smeared consistency with indistinguishable structure, grayish-red in color and 10*3 cm in size. The blood in the lumen of the 4th ventricle, 3rd ventricle and lateral ventricles. | Perivascular and pericellular edema, fresh hemorrhage in the brain matter. | Intracerebral hemorrhage with rupture into the ventricular system. | The cerebral edema with occlusion of the cerebellar tonsils into the greater occipital foramen. | 3.0 |
| 15 | 558/20 | 69 | F | Cerebral cortex. | In the area of the frontal lobe of the left cerebral hemisphere there was an 8x4x3 cm hemorrhage. | Perivascular and pericellular edema, fresh hemorrhage in the brain matter. | Intracerebral hemorrhage in the left frontal lobe. Massive subarachnoid hemorrhage. | The cerebral edema with occlusion of the cerebellar tonsils into the greater occipital foramen. | 3.0 |
| 16 | 601/20 | 65 |  | Cerebral cortex. | In the area of the frontal lobe of the right cerebral hemisphere, a hemorrhage area sized 7x5x3 cm. | Perivascular and pericellular edema, fresh hemorrhage in the brain matter. | Intracerebral hemorrhage in the right frontal lobe. | The cerebral edema with occlusion of the cerebellar tonsils into the greater occipital foramen. | 3.0 |
| 17 | 60/20 | 52 | F | Cerebral cortex. | In the area of the parietal lobe of the right hemisphere there is a 6x4x4 cm cavity filled with a blood clot, with a defect in the wall, through which blood flows into the lumen of the ventricles. | Perivascular and pericellular edema, fresh hemorrhage in the brain matter. | Intracerebral hemorrhage with formation of intracerebral hematoma in the right hemisphere and rupture into the ventricular system of the brain. | The cerebral edema with occlusion of the cerebellar tonsils into the greater occipital foramen. | 3.0 |
| 18 | 26/19 | 75 | M | Cerebral cortex. | In the basal nuclei of the right hemisphere there is a cavity of 6x5x4 cm filled with blood clot, there is a defect in the wall, through which blood enters the ventricular lumen. | Perivascular and pericellular edema, fresh hemorrhage in the brain matter. | Intracerebral hemorrhage in the right hemisphere with formation of intracerebral hematoma. | The cerebral edema with occlusion of the cerebellar tonsils into the greater occipital foramen. | 2.4 |
| 19 | 38/19 | 78 | F | Cerebral cortex. | In the area of the temporoparietal lobe of the left hemisphere there is a cavity of 5x7x5 cm filled with blood clot, there is a defect in the wall, through which blood flows into the lumen of the ventricles. | Perivascular and pericellular edema, fresh hemorrhage in the brain matter. | In the area of the temporo-parietal lobe of the left hemisphere there is a cavity of 5x7x5 cm filled with blood clot, there is a defect in the wall, through which blood flows into the lumen of the ventricles. | The cerebral edema with occlusion of the cerebellar tonsils into the greater occipital foramen | 2.1 |
| 20 | 27/19 | 79 | F | Cerebral cortex. | The lumen of the fourth and lateral ventricles contains 150 ml of blood. | Perivascular and pericellular edema, fresh hemorrhage in the brain matter. | Spontaneous recurrent intracerebral hemorrhage of multiple localization. | The cerebral edema with occlusion of the cerebellar tonsils into the greater occipital foramen. | 2.5 |
| 21 | 53/19 | 61 | F | Cerebral cortex. | In the right occipital lobe, a softening foci of smeared consistency with indistinguishable structure, grayish-red and 5*5*6 in size, blood in the lumen of the left lateral ventricle | Perivascular and pericellular edema, fresh hemorrhage in the brain matter. | Parenchymal and ventricular hemorrhage with formation of intracerebral hematoma in the left cerebral hemisphere with blood rupture into the ventricles. | The cerebral edema with occlusion of the cerebellar tonsils into the greater occipital foramen. | 3.0 |
| 22 | 56/19 | 76 | F | Cerebral cortex. | The left occipital lobe has a focal hemorrhage sized 4*3*8, with blood in the lumen of the left lateral ventricle. | Perivascular and pericellular edema, fresh hemorrhage in the brain matter. | Parenchymal and ventricular hemorrhage in the left cerebral hemisphere with formation of subdural and intracerebral hematomas with blood rupture into ventricles and subarachnoid space. | The cerebral edema with occlusion of the cerebellar tonsils into the greater occipital foramen. | 3.0 |
| 23 | 346/20 | 76 | M | Cerebral cortex. | In the area of the parietal lobe of the left hemisphere there is a 5x5x6 cm cavity filled with blood clot, there is a defect in the wall, through which blood flows into the lumen of the ventricles. | Perivascular and pericellular edema, fresh hemorrhage in the brain matter. | Parenchymatous ventricular hemorrhage with formation of intracerebral hematoma in the left cerebral hemisphere with blood rupture into the brain ventricles, motor-sensory aphasia, right hemiplegia. | The cerebral edema with occlusion of the cerebellar tonsils into the greater occipital foramen. | 3.0 |
| 24 | 361/20 | 71 | M | Cerebral cortex. | In the area of basal nuclei there was a softening foci of smeary consistency with indistinguishable structure, grayish-red in color and 3*4*3 in size, there was blood in the ventricular lumen. | Perivascular and pericellular edema, fresh hemorrhage in the brain matter. | Consequences of intracerebral hemorrhage in the left middle cerebral artery basin with formation of intracerebral hematoma in the basal ganglia with rupture into the brain ventricles. | The cerebral edema with occlusion of the cerebellar tonsils into the greater occipital foramen. | 3.0 |
| 25 | 160/20 | 75 | F | Cerebral cortex. | Hemorrhage in the basal nuclei of the left hemisphere with blood rupture into the lateral ventricles of the brain. | Perivascular and pericellular edema, fresh hemorrhage in the brain matter. | Parenchymatous ventricular hemorrhage in the left cerebral hemisphere with formation of intracerebral hematoma with blood rupture into the ventricles. | The cerebral edema with occlusion of the cerebellar tonsils into the greater occipital foramen. | 3.0 |
| 26 | 76/20 | 55 | M | Cerebral cortex. | There is blood in the lumen of the 4 ventricles and lateral ventricles. In the left posterior communicating artery, a saccular aneurysm with a diameter of 0.5 cm with a rupture. The wall of the aneurysm with a layered thrombus of a gray-cross color. | Perivascular and pericellular edema, fresh hemorrhage in the brain matter.  **Immunohistochemical staining of cerebral cortex tissues revealed Lyve1/Prox-1 positive elements.** | There was blood in the lumen of the 4th ventricle and lateral ventricles. In the left posterior connecting artery there is a 0.5 cm diameter sac-shaped aneurysm with rupture. The aneurysm wall was with laminated gray-cross-colored thrombi. | The cerebral edema with occlusion of the cerebellar tonsils into the greater occipital foramen. | 2.5 |
| 27 | 160/20 | 52 | M | Cerebral cortex. | The subdural space on the right side contains a small amount of blood, infiltrating the cerebral cortex. | Perivascular and pericellular edema, fresh hemorrhage in the brain matter. | Spontaneous subarachnoid hemorrhage with formation of subdural hematoma in the right cerebral hemisphere. | The cerebral edema with occlusion of the cerebellar tonsils into the greater occipital foramen. | 2.1 |
| 28 | 62/20 | 61 | M | Cerebral cortex. | Hemorrhage in the parietal lobe of the right cerebral hemisphere sized 5x4 cm. | Perivascular and pericellular edema, fresh hemorrhage in the brain matter. | Parenchymatous hemorrhage with formation of intracerebral hematoma in the right cerebral hemisphere. | The cerebral edema with occlusion of the cerebellar tonsils into the greater occipital foramen. | 2.5 |
| 29 | 192/20 | 67 | F | Cerebral cortex. | There was a 100 ml hemorrhage in the left brainstem nuclei with rupture into the fourth and lateral ventricles. | Perivascular and pericellular edema, fresh hemorrhage in the brain matter. | Parenchymal-ventricular hemorrhage in the left hemisphere of the brain with the formation of an intracerebral hematoma and penetration of blood into the ventricles. | The cerebral edema with occlusion of the cerebellar tonsils into the greater occipital foramen. | 2.5 |
| 30 | 13/20 | 52 | M | Cerebral cortex. | In the right cerebral hemisphere from the frontal to the parietal lobe there was a hemorrhage of 9x4 cm in size. There was blood in the lumen of the 4th ventricle, 3rd ventricle and lateral ventricles. | Perivascular and pericellular edema, fresh hemorrhage in the brain matter. | Intracerebral hemorrhage with a penetration into the ventricular system. | The cerebral edema with occlusion of the cerebellar tonsils into the greater occipital foramen. | 3.0 |
| 31 | 245/20 | 65 | F | Cerebral cortex. | In the area of the temporal lobe of the left cerebral hemisphere, there was an area of hemorrhage measuring 8x4x3 cm. | Perivascular and pericellular edema, fresh hemorrhage in the brain matter. | Intracerebral hemorrhage in the left frontal lobe. Massive subarachnoid hemorrhage. | The cerebral edema with occlusion of the cerebellar tonsils into the greater occipital foramen. | 3.0 |
| 32 | 122/20 | 64 | F | Cerebral cortex. | In the area of the frontal lobe of the right cerebral hemisphere there was a hemorrhage measuring 6x4x3 cm. | Perivascular and pericellular edema, fresh hemorrhage in the brain matter. | In the area of the frontal lobe of the right cerebral hemisphere there was a hemorrhage measuring 6x4x3 cm. | The cerebral edema with occlusion of the cerebellar tonsils into the greater occipital foramen. | 3.0 |
| 33 | 53/20 | 74 | M | Cerebral cortex. | In the area of the occipitoparietal lobe of the left hemisphere, a cavity measuring 6x7 cm filled with a blood clot, there is a defect in the wall through which blood flows into the lumen of the ventricles. | Perivascular and pericellular edema, fresh hemorrhage in the brain matter. | Parenchymatous ventricular hemorrhage with formation of intracerebral hematoma in the left cerebral hemisphere with blood rupture into the brain ventricles, motor-sensory aphasia, right hemiplegia. | The cerebral edema with occlusion of the cerebellar tonsils into the greater occipital foramen. | 3.0 |
| 34 | 352/20 | 71 | F | Cerebral cortex. | In the basal nuclei there was a softening foci of smeared consistency with indistinguishable structure, grayish-red color and size 4*4*4, there was blood in the ventricular lumen. | Perivascular and pericellular edema, fresh hemorrhage in the brain matter. | Consequences of intracerebral hemorrhage in the left middle cerebral artery basin with formation of intracerebral hematoma in the basal ganglia with rupture into the brain ventricles. | The cerebral edema with occlusion of the cerebellar tonsils into the greater occipital foramen. | 2.1 |

Table S2. The control group of patients, who died from the congestive heart failure with the development of pulmonary edema

| **N** | **HSB Sample no** | **Age** | **Sex** | **Structure** | **Neuropathology Macroscopic description** | **Neuropathology Microscopic description** | **Clinical diagnosis** | **Cause of death** | **Postmortem time, h** |
| --- | --- | --- | --- | --- | --- | --- | --- | --- | --- |
| 1 | 382/20 | 65 | M | Cerebral cortex. | The brain has elastic consistency to the touch, whitish-grayish color on sections, surfaces of sections are smooth, shiny, structure of cerebral nuclei and brain stem preserved, ependyma of ventricles is smooth, shiny, traces of transparent light-yellow fluid in their lumens. | Normal structure of cerebral tissues, very mild perivascular edema. | Postinfarction cardiosclerosis. Stenotic atherosclerosis of the coronary arteries of the heart (grade 3; stage II; stenosis up to 70%). | Congestive heart failure with the development of pulmonary edema. | 2.5 |
| 2 | 381/20 | 79 | M | Cerebral cortex. | The brain has elastic consistency to the touch, whitish-grayish color on sections, surfaces of sections are smooth, shiny, structure of cerebral nuclei and brain stem preserved, ependyma of ventricles is smooth, shiny, traces of transparent light-yellow fluid in their lumens. | Normal structure of cerebral tissues, very mild perivascular edema. | Postinfarction cardiosclerosis. Stenotic atherosclerosis of coronary arteries of the heart (degree 3; stage III; stenosis up to 60%). | Congestive heart failure with the development of pulmonary edema. | 2.0 |
| 3 | 879/20 | 61 | F | Cerebral cortex. | The brain has elastic consistency to the touch, whitish-grayish color on sections, surfaces of sections are smooth, shiny, structure of cerebral nuclei and brain stem preserved, ependyma of ventricles is smooth, shiny, traces of transparent light-yellow fluid in their lumens. | Normal structure of cerebral tissues, very mild perivascular edema.  **Immunohistochemical staining of cerebral cortex tissues revealed Lyve1/Prox-1 positive vessels and elements.** | Postinfarction cardiosclerosis. Stenotic atherosclerosis of the coronary arteries of the heart (grade 3; stage II; stenosis up to 70%). | Congestive heart failure with the development of pulmonary edema. | 2.0 |
| 4 | 364/20 | 87 | F | Cerebral cortex. | The brain has elastic consistency to the touch, whitish-grayish color on sections, surfaces of sections are smooth, shiny, structure of cerebral nuclei and brain stem preserved, ependyma of ventricles is smooth, shiny, traces of transparent light-yellow fluid in their lumens. | Normal structure of cerebral tissues, very mild perivascular edema. | Postinfarction cardiosclerosis. Stenotic atherosclerosis of coronary arteries of the heart (degree 3; stage III; stenosis up to 65%). | Congestive heart failure with the development of pulmonary edema. | 2.1 |
| 5 | 51/20 | 59 | F | Cerebral cortex. | The brain has elastic consistency to the touch, whitish-grayish color on sections, surfaces of sections are smooth, shiny, structure of cerebral nuclei and brain stem preserved, ependyma of ventricles is smooth, shiny, traces of transparent light-yellow fluid in their lumens. | Normal structure of cerebral tissues, very mild perivascular edema. | Postinfarction cardiosclerosis. Stenotic atherosclerosis of the coronary arteries of the heart (degree 3; stage IV; stenosis up to 70%). | congestive heart failure with the development of pulmonary edema. | 2.0 |
| 6 | 223/20 | 74 | M | Cerebral cortex. | The brain has elastic consistency to the touch, whitish-grayish color on sections, surfaces of sections are smooth, shiny, structure of cerebral nuclei and brain stem preserved, ependyma of ventricles is smooth, shiny, traces of transparent light-yellow fluid in their lumens. | Normal structure of cerebral tissues, very mild perivascular edema.  **Immunohistochemical staining of cerebral cortex tissues revealed Lyve1/Prox-1 positive vessels and elements.** | Postinfarction cardiosclerosis. Stenotic atherosclerosis of coronary arteries of the heart (degree 3; stage III; stenosis up to 65%). | Congestive heart failure with the development of pulmonary edema. | 2.5 |
| 7 | 497/20 | 76 | M | Cerebral cortex. | The brain has elastic consistency to the touch, whitish-grayish color on sections, surfaces of sections are smooth, shiny, structure of cerebral nuclei and brain stem preserved, ependyma of ventricles is smooth, shiny, traces of transparent light-yellow fluid in their lumens. | Normal structure of cerebral tissues, very mild perivascular edema.  **Immunohistochemical staining of cerebral cortex tissues revealed Lyve1/Prox-1 positive vessels and elements.** | Postinfarction cardiosclerosis. Stenotic atherosclerosis of coronary arteries of the heart (degree 3; stage IV; stenosis up to 70%). | Congestive heart failure with the development of pulmonary edema. | 3.0 |
| 8 | 756/21 | 61 | M | Cerebral cortex. | The brain has elastic consistency to the touch, whitish-grayish color on sections, surfaces of sections are smooth, shiny, structure of cerebral nuclei and brain stem preserved, ependyma of ventricles is smooth, shiny, traces of transparent light-yellow fluid in their lumens. | Normal structure of cerebral tissues, very mild perivascular edema. | Liver cirrhosis. | Bleeding from esophageal veins. Posthemorrhagic anemia. Congestive heart failure with the development of pulmonary edema. | 2.5 |

Table S3. The quantitative analysis of PVS, CD31, CD68 and Lyve-1 positive structures

| Groups/Number of brain samples | **The diameter of CD31-positive blood vessels (μm)** | The number of CD68-positive macrophages in PVS | The diameter of Lyve1-positive structures (μm) | The number of Lyve1 structures | **The size of PVS, μm** |
| --- | --- | --- | --- | --- | --- |
| **Мe**  (Min-Max) Q [25-75] | | | | | |
| **The Lyve+group with IVH, n=8** | 20  (11 - 80)  [14-30] | 2,5  (0-8)  [0,25-4] | 10  (10-90)  [10-20] | 3  (1-6)  [2-4,5] | 18  (10-42)  [15-20] |
| **The Lyve- control group, n=3** | 19  (10-41)  [14-22] | 1  (0-3)  [0-3] | 10  (0-20)  [10-12] | 1  (0-2) | 2  (0-4)  [0-4] |
| **Mann-Whitney criterion** | Z= -1,34  p=0,18 | Z= -2,26  p=0,023 | Z= -1,75  p=0,07 | Z=-5,08  p=3,7*10^-7^ | Z= -6,7  p=2,03* 10^-11^ |
| **The Lyve- group with IVH, n=26** | 17  (5-22)  [9-21] | 0  (0-1)  [0-1] | - | - | 9  (3-17)  [6-13] |
| **The Lyve+ control group, n=5** | 12  (1-10)  [10-18] | 0  (0-1)  [0-1] | - | - | 0  (0-1)  [0-0.5] |
| **Mann-Whitney criterion** | Z= -0,41  p=0,67 | Z= -0,59  p=0,55 | Z=0  p=1 | Z=0  p=1 | Z= -6,05  p=1,4*10^-9^ |

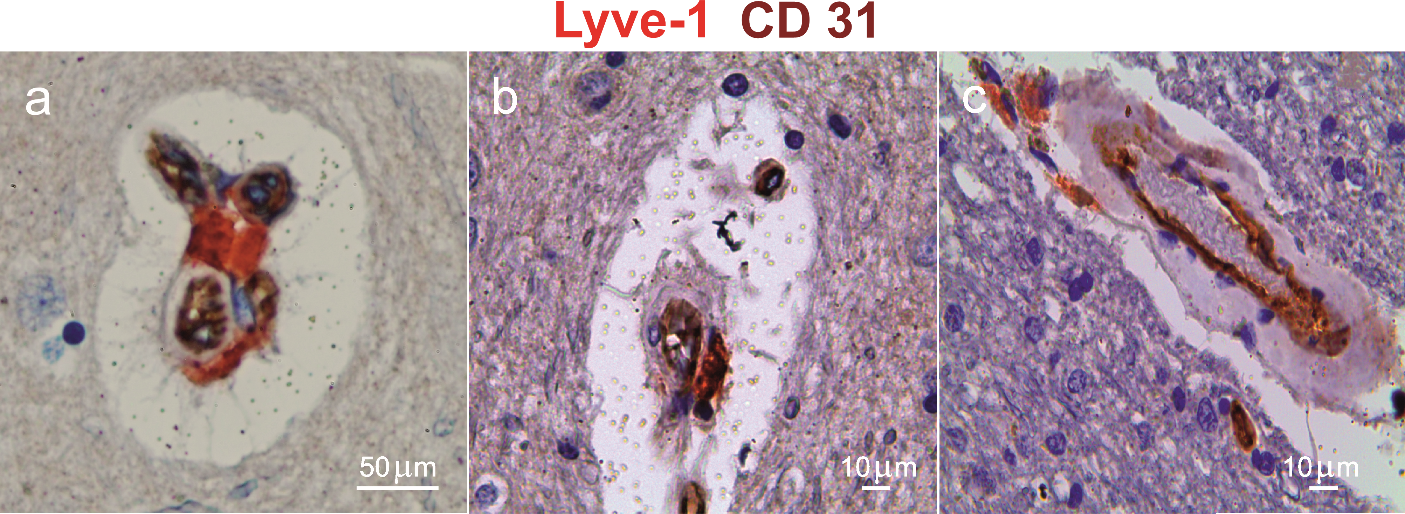

**Fig. S1. Representative images of Lyve-1-expressing structures close to CD31-expressing capillary (a), venula (b) and arteriole (c).** 10 fields from one brain were analyzed, n=3 for the control group and n=8 for the IVH group.

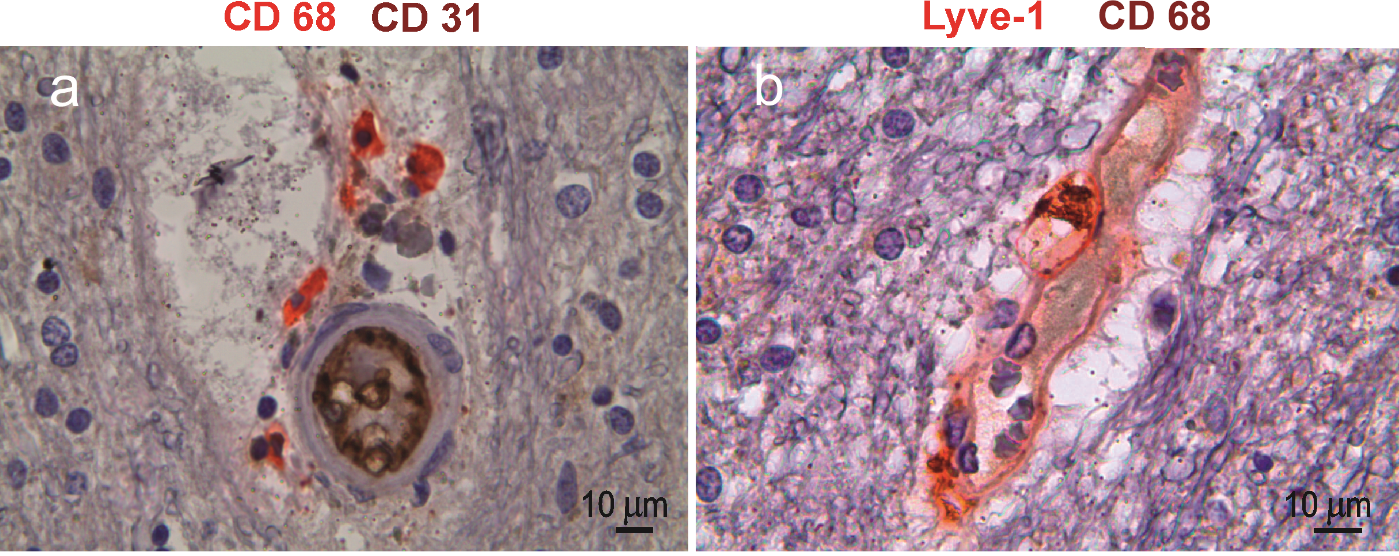

Fig. S2. Representative images of CD68-expressing cells in the area of vasogenic edema (a) and inside of Lyve-1-experssing structures (b) in the human brain with IVH. 10 fields from one brain were analyzed, n=8 for the IVH group.

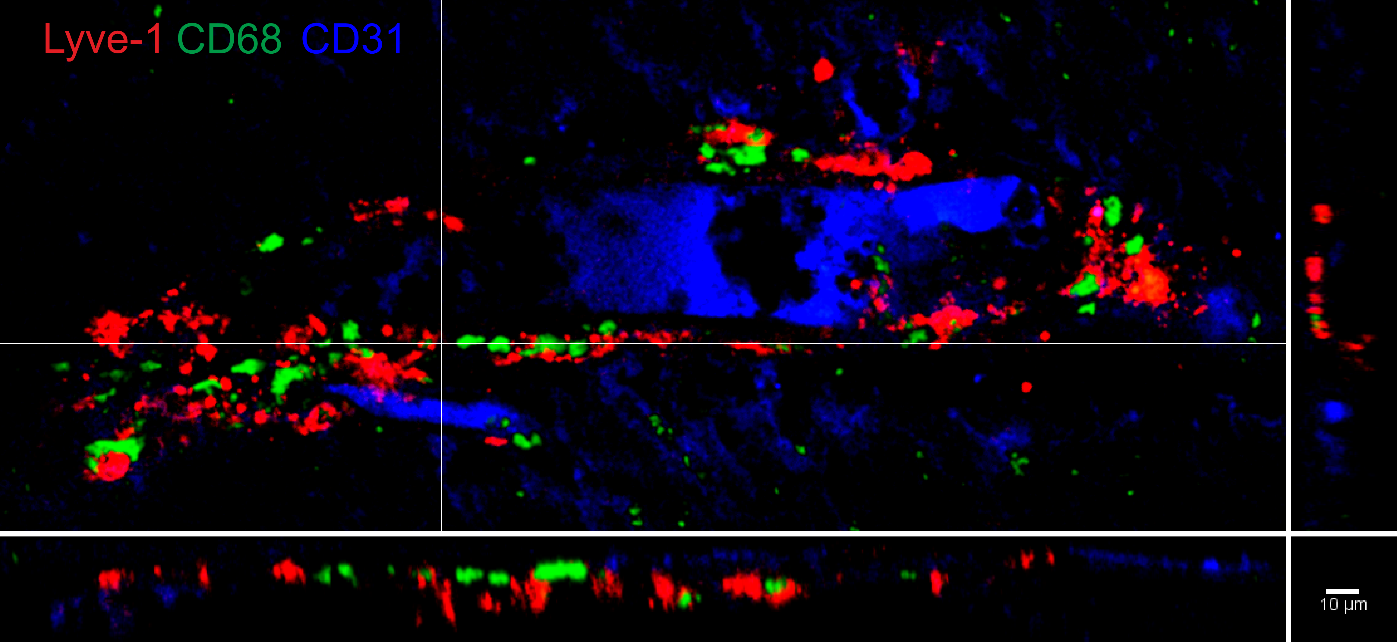

**Fig. S3. Representative image of Lyve-1+, СD68+ and CD31+ cells in the human brain with IVH.** 10 fields from one brain were analyzed, n=9 for the IVH group.

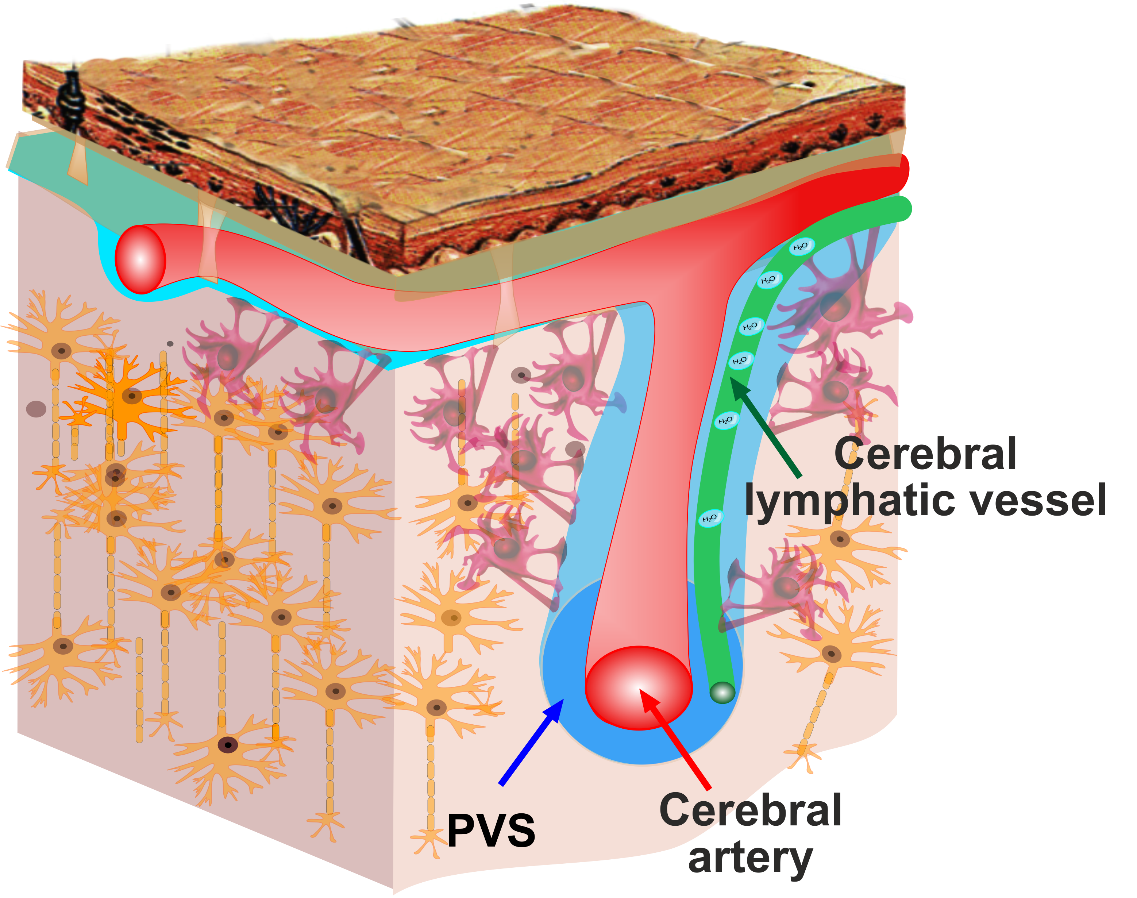

**Fig. S4. The schematic representation of LVs in the human brain and a model of brain lymphatic drainage.** The LVs are presented along the cerebral blood vessels, express Lyve-1/Prox-1 and have valves, a single endothelial layer, undulating shape of lymphatic endothelial cells. The pulsation of the cerebral arteries is the driving force behind the movement of fluids from the perivascular spaces to the extracellular space. The cerebral lymphatic vessels direct the brain fluids from the perivascular spaces to the subarachnoid space.

**Table S4. The general characteristics of the initial lymphatics and precollectors**

| **Characteristics** | **References** | **Methods of detection of LVs characteristics in this study** |
| --- | --- | --- |
| Lyve1/Prox1 expression. | Nature. 523, 337–341 (2015)  J. Exp. Med. 212, 991–999 (2015)  Nature. 572, 62–66 (2019) | Confocal colocolozation analysis |
| Initial LVs have undulating shape of lymphatic endothelial cells.  Precollectors are defined as LVs composed of a single endothelial layer but also having secondary valves to prevent backflow into initial lymphatics. | Compr Physiol. 9(1): 207–299 (2018).  J Cell Biol. 193(4), 607-618 (2011) | Confocal analysis |
| Initial LVs have single endothelial layer with an indistinct basal lamina and an absence of muscle layer. | Arch. Histol. Cytol. 53, 95-105 (1990) | Electronic microscopy |

Supplementary Text

**Modeling of brain drainage**

The pathways of cerebral fluids that ensure the timely removal of metabolic waste has been discussed for a long time and intensively. Currently, the glymphatic hypothesis proposed in 2012 (14,15) is most discussible. Some of its assumptions have been confirmed experimentally and can be regarded as a proven facts. These include the presence of directional cerebrospinal fluid (CSF) current in the perivascular spaces (PVS) of the pial arteries and the important role of vascular pulsation (16-18).

At the same time, there is no common agreement on another their key parts. Specifically, the work (19) did not reveal a directed flow in the parenchyma, and the mathematical calculation of the characteristics of peristaltic transport shows that it is too weak to provide the necessary flow (20). There are also contradicting views on the calculated pressure gradient (21,22).

Note that another schemes of fluid flows, different from the glymphatic hypothesis, have been proposed. In works by Weller et al. (23,24) it was argued that the outflow of fluid from the parenchyma occurs through the intercellular space of the vascular wall, as shown in Fig. S5, panel A.

In (25) the glymphatic hypothesis is compared with an alternative mechanism based on diffusion, as shown in Figure S5, panels B and C, respectively. The major role of diffusion is also argued in (19,26). To summarize, currently there is still no common agreement about the pathways of drainage of the brain parenchyma.

At the same time, the works (16, 17) convincingly demonstrated the presence of a fluid flow directed towards the parenchyma in PVS of arterial vessels, and it can be seen from the visualized particle trajectories that their movement is affected by pulsations, but is not created by them. Thus, inconsistencies still exist both in the general picture of the movement of fluids and in the specific mechanism of action of pulsations.

The presence of the lymphatic vessels (LVs) as dedicated channels for the drainage of fluid through the vascular wall allows a return to the ideas of Weller et al. (23, 24) at a new level, replacing the outflow of fluid through the intercellular space of the smooth muscle cells of the vessel for its movement along the found channel.

Note that it is critically important for drainage efficiency that the drainage path should have a low resistance to flow, since the pressure gradient between the CSF and the lymph vessels of the meninges is small (22). The detected in our work LVs have the diameter about 10-20 microns. If we assume that the transverse size of the intercellular space gaps inside the vascular wall is 0.2 microns (27), then the calculation using the Hagen-Poiseuille flow formulas (28) in a round tube and a thin flat channel shows that the fluid flow through LVs will be 20-500 more than through a narrow slit assuming the same cross-sectional area and pressure gradient.

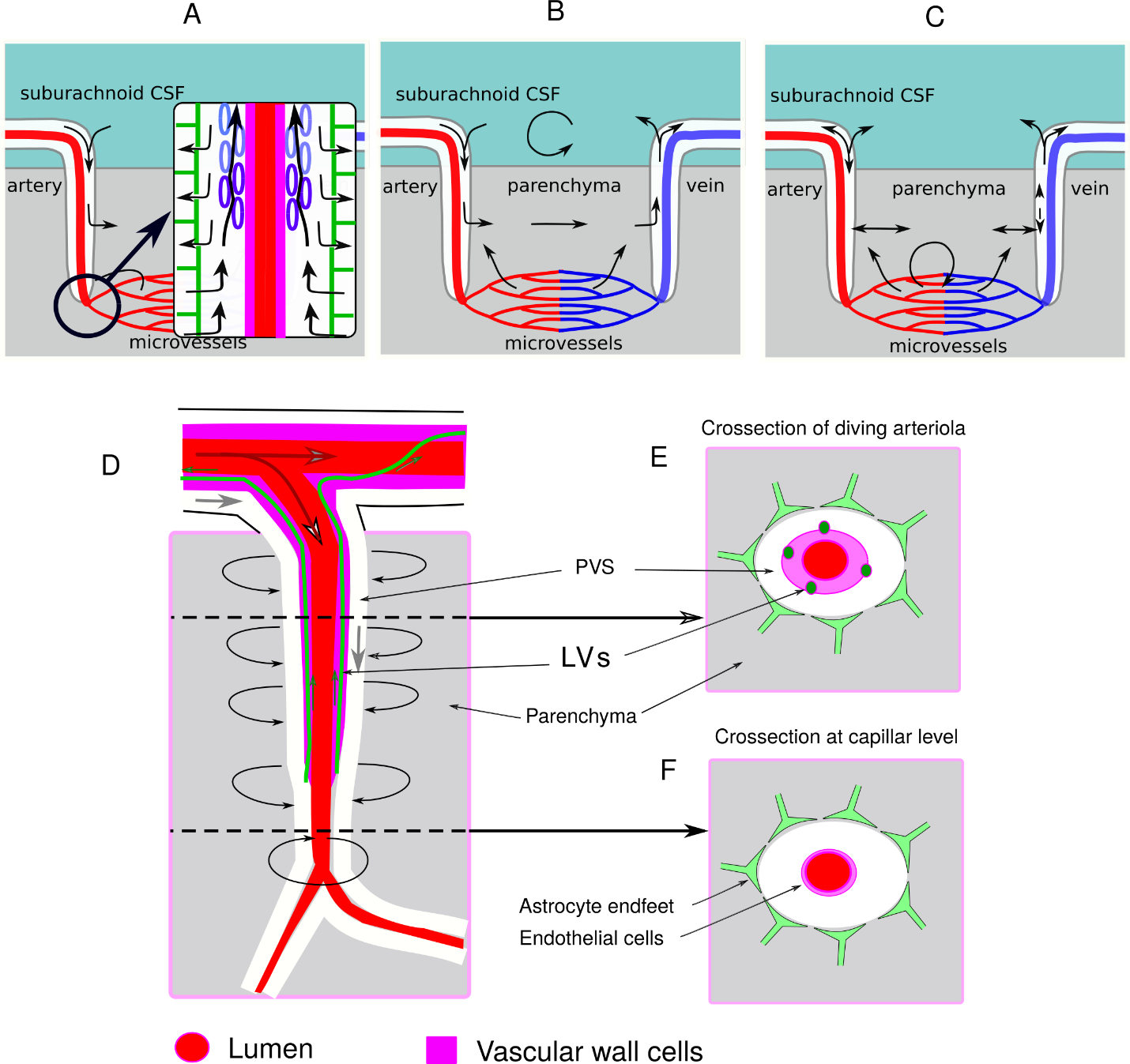

**Fig. S5 – Schemes of possible fluid motion to and from the brain parenchyma.** A- according to (30,31); B,C – according to (32); D,E,F – based on LVs existence.

Based on this fact, the diagram of the movement of fluids and drainage of the parenchyma is shown in Fig. S5 D, E, F, where the pulsations of the arterial vessel play the key role, pushing the CSF both from the PVS into the parenchyma (Prn) and back. Thus, bulk flow through the parenchyma is absent or at least very small compared to the flow in the PVS, and the transfer of metabolic waste from the brain parenchyma to PVS is accelerated by the process of the combined action of pulsatile transfer (advection) and diffusion, as discussed earlier in (29).

Quantitative estimation of the contribution of such a pulsating flow to the drainage process requires a dedicated model study, which is beyond the scope of this work and will be published separately. Here we will illustrate just the principle playability of the proposed mechanism using a simple physical model consisting of 9 compartments, as shown in Fig. S6 A. Each of 9 compartments is described by the same set of equations as shown in Fig. S6 B. The desired structure of the simulated system is set by both local parameters for each compartment, and its connections with neighboring elements of the model. In particular, effective compliance for PVS elements was the explicit function of time, which made it possible to simulate pulsations.

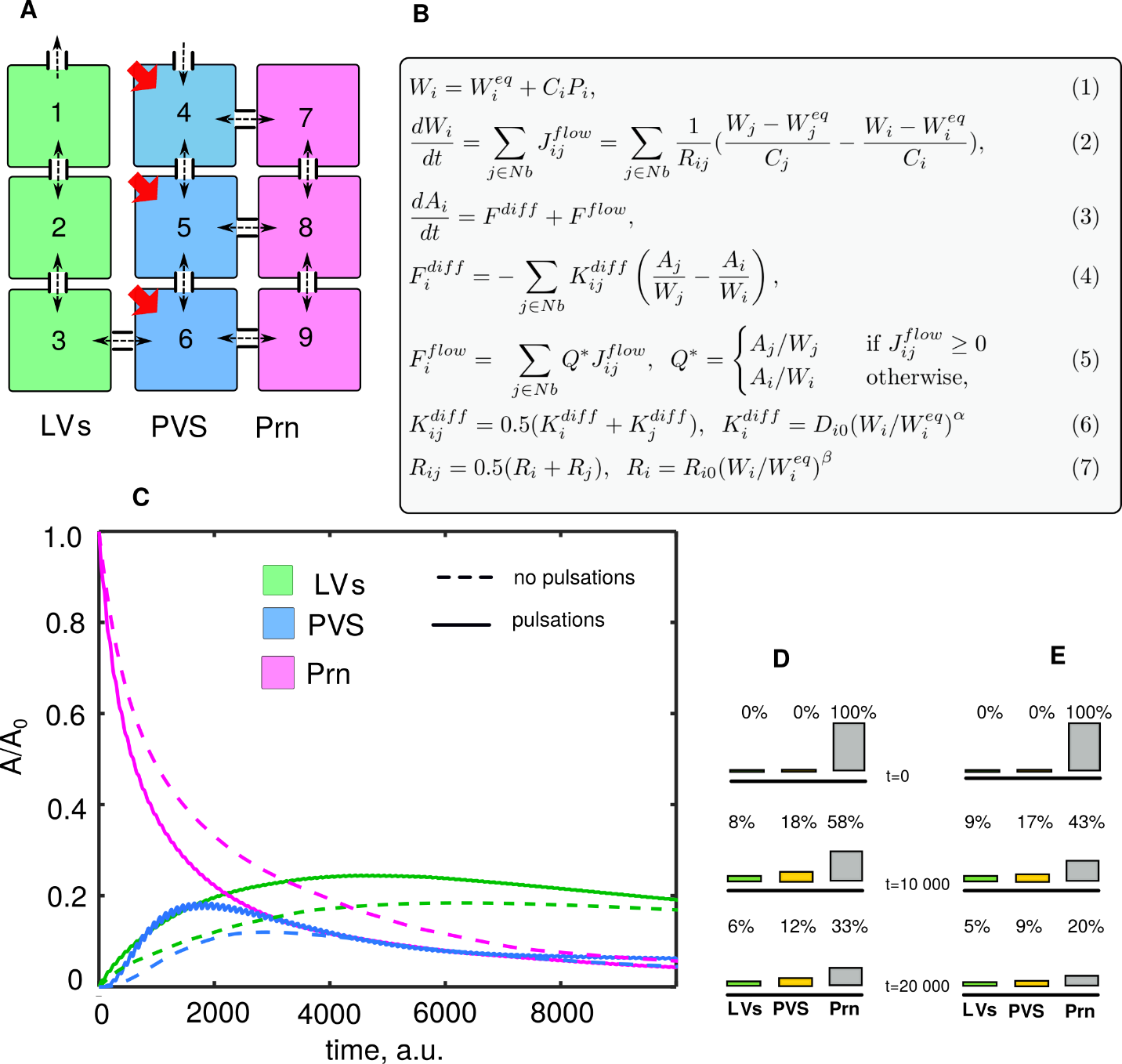

**Fig. S6** **- Modeling of pulsation assisted drainage.** A:The 9 compartments of the model; B: The equation for each of the model compartments; C: The modeled washout of a substance, initially loaded in parenchyma only (1.0 at t=0), without (dashel lines) and with (solid lines) arterial pulsations. D and E: The bar diagrams showing the spreading of model substance with time between the lymphatic vessels (LVs), the perivascular space (PVS), and the brain parenchyma (Prn).

In the course of simulations, the initial conditions were set such that all substance *A* (which represents a "metabolic waste") is loaded in the parenchyma (compartments 7,8,9). To take into account the hindered coupling between the PVS and the parenchyma, both the inverse hydrodynamic resistance *1/Rij* and the diffusion rate *K^diff^_ij_* at junctions of elements 4-7, 5-8, and 6-9 were reduced by a factor of 10 in comparison with coupling between elements of the same type, such as 1-2 , 2-3, 3-6, 4-5, 5-6, 7-8, 8-9. In Fig. 6S C,D, and E we compared two cases: when the compliance of PVS elements 4,5,6 was constant and when it periodically decreased, which mimicked the effect of arterial pulsation.

One can observe the presence of pulsations is indeed capable of accelerating the washing out of metabolic debris from the parenchyma under conditions when the main flow during drainage does not pass through the parenchyma.
